# The two-component microbial system of the black soldier fly larvae (BSFL) gut: a plastic microbiota in the midgut, but a stable one in the hindgut

**DOI:** 10.64898/2026.09.26.754629

**Authors:** Maurielle Eke, Kevin Tougeron, Manon Martin, Jérôme Ambroise, Bertrand Bearzatto, Jean-Luc Gala, Leonard S. Ngamo Tinkeu, Thierry Hance, François Renoz

**Author notes:** Address correspondence to François Renoz.

## Abstract

Due to their highly polyphagous capacities, black soldier fly (*Hermetia illucens*) larvae (BSFL) are increasingly valued for their ability to convert organic waste into valuable biomass that can be used for a variety of purposes. These remarkable digestive capabilities are highly dependent on an extremely plastic gut microbiota. However, the distribution and functioning of bacterial communities in the various gut compartments - particularly in the hindgut - remain little understood. In this study, we used a metabarcoding approach based on 16S gene sequencing to investigate the effect of three carbohydrate-rich diets with distinct molecular compositions on the functional diversity of the BSFL gut microbiota. Our results showed that the midgut harbors a highly substrate-sensitive microbiota, with a high abundance of *Actinomyces* spp., regardless of the substrate. A bacterial diversity oriented toward fatty acid biosynthesis pathways is promoted by starch-rich environment, whereas a lignocellulosic substrate fosters a midgut microbiota dominated by *Paenibacillus* spp. In contrast, the hindgut exhibits a distinctly stable and homogeneous bacterial composition dominated by *Dysgonomonas* spp. Overall, our results provide clear evidence of a two-compartment microbial system, in which the midgut primarily serves as a substrate-adaptive primary degradation chamber, while the hindgut functions as a stable terminal compartment for the final processing of residual substrates and the recycling of nutrients. These findings contribute to our understanding of the functional diversity of the bacterial microbiota along the BSFL digestive tract, which is a key factor in explaining this insect’s remarkable polyphagous behavior and optimizing its use for industrial purposes.

**Importance:** By converting various types of organic waste into protein- and lipid-rich larvae, black soldier fly (BSFL) larvae act as a flexible bio-factory driven by their digestive tract. Yet, the functional diversity of the BSFL gut microbiota remains poorly characterized across the compartments of the digestive tract, with the hindgut still a black box. Grasping these aspects is essential to elucidate how this bio-factory functions and to guide the production of larvae tailored to specific industrial needs. By examining how different carbohydrate-rich diets affect this functional diversity, our research provides clear evidence for a two-compartment microbial system: the midgut functioning as a substrate-adaptive primary degradation chamber, and the hindgut functioning as a stable terminal compartment that processes residual substrate and recycles nutrients, with the genus *Dysgonomonas* acting as a key taxon. Furthermore, our research identifies pivotal bacterial genera (including *Actinomyces* and *Paenibacillus*) supporting BSFL polyphagy under carbohydrate-rich diets.

## Introduction

The increase in food demand, driven by global population growth, generates huge amounts of organic waste (food waste, livestock byproducts, etc.) that have adverse environmental, social, and economic impacts (1). One strategy for managing this waste involves integrating it into a sustainable circular economy to transform it into valuable biomass (2). In this context, insect-based bioconversion is an emerging research topic and a rapidly expanding commercial venture (3). Larval biomass derived from the conversion of organic waste can be used for multiple purposes, including as a source of protein and lipids for animal and human food, pharmaceuticals, biofuels, lubricants, biogas, and fertilizers (4–10). In recent years, black soldier fly larvae (BSFL, *Hermetia illucens*) have emerged as the most efficient known bioconversion organisms, capable of growing on an incredible variety of organic substrates, ranging from fresh fruits and vegetables to lignocellulosic materials and antinutritional-rich residues (11). BSFL operate as flexible bio-factories enabling the conversion of organic waste into protein- and lipid-rich larvae that can be used for a variety of purposes (6, 12–15). Understanding how these bio-factories work and determining how to optimize them to produce larvae whose chemical profile meets market demands is a major challenge.

In recent years, it has become increasingly evident that the polyphagous capacity of BSFL is not solely attributable to their own enzymatic system, but is primarily regulated by a highly plastic gut microbiota, whose composition is tailored to the chemical nature of the ingested substrate (16–19). Culture-based and metabarcoding approaches have provided a detailed picture of the BSFL gut microbiota on various substrates and under different conditions (20). While the nature of the substrate is a key factor in shaping the BSFL microbiota, other factors may also come into play, including the insect’s developmental stage, rearing temperature, exposure to toxic compounds, and the host genome. Although certain bacterial genera appear to be pervasive in the BSFL gut (*e.g., Morganella*, *Providencia*, *Dysgonomonas*, *Ignatzschineria*, *Proteus*, *Bacillus*, and *Klebsiella*), the existence of a core microbiota in the strict sense remains uncertain. Furthermore, despite the numerous studies devoted to the BSFL microbiota, one aspect that remains elusive is the specific role that these gut-associated bacteria may play in digestion. This gap is largely attributed to the fact that the BSFL bacterial microbiota has rarely been studied with due consideration for the anatomical complexity of this insect’s digestive system: most metabarcoding studies have used the entire digestive tract as biological material for their analyses, without considering gut compartmentalization (20). Only a few recent studies have advanced our knowledge in this area by demonstrating that the bacterial communities of BSFL are organized according to the different compartments of the digestive tract (17, 21–24). As in all insects, the digestive tract of *H. illucens* larvae is organized into distinct morphological and functional regions: the foregut, midgut, and hindgut (25). The midgut of BSFL is subdivided into anterior (pH ∼6), middle (pH ∼2), and posterior (pH ∼8.5) regions (21, 26), each harboring its own resident microbiota (17, 22–24). While bacterial diversity specifically associated with the midgut is now beginning to be considered, that associated with the hindgut remains largely neglected. It is now timely to go further and identify the role of associated bacteria in the digestion of organic waste across the different compartments of the digestive tract, to better understand the evolutionary ecology of *H. illucens*, explain its polyphagous behavior, and optimize its use in practical applications.

In a previous study, we demonstrated that different high-carbohydrate diets can be used to customize the protein and lipid content of BSFL (27). Specifically, we found that a diet consisting of forage produces protein-rich larvae, while a diet consisting entirely of potato scraps increases the fatty acid content of BSFL. These results are meaningful because they demonstrate that carbohydrate-based diets with differing macromolecular compositions can promote the production of BSFL with markedly different value-added properties. The bioconversion profiles we previously identified provide a solid foundation for studying the underlying mechanisms, particularly the role of the BSFL gut microbiota.

In this study, we examined the effect of different high-carbohydrate diets on the functional diversity of the gut microbiota of BSFL. This study is part of ongoing efforts to optimize the use of organic waste for the production of high-quality larvae for poultry farming (27, 28). We reared BSFL on three distinct carbohydrate-rich diets (diet A: 100% fruit/vegetable waste; diet B: 100% potato scraps waste; diet C: 100% forage) and used a metabarcoding approach to determine how these three diets shape bacterial communities in the midgut *versus* the hindgut of BSFL. To shed light on the functional role of gut microbiota members, we correlated these results with the substrate and larval composition obtained previously (27), and used PICRUSt2 to infer the putative role of bacteria associated with BSFL in the digestion of carbohydrate-rich diets. We hypothesized that (i) gut bacterial communities are structured according to anatomical compartment (midgut *versus* hindgut); (ii) the structure of the gut microbial communities shifts according to the type of diet, with distinct bacterial taxa enriched in response to specific carbohydrate sources; and (iii) these compositional changes are reflected in predicted metabolic profiles. Overall, our results provide clear evidence of a two-compartment microbial system, in which the midgut primarily serves as a substrate-adaptive primary degradation chamber, while the hindgut processes residual substates and recycles nutrients, with *Dysgonomonas* acting as a key taxon.

## Results

### 1. The gut compartment is the main determinant of microbial community composition

The impact of the three carbohydrate-rich diets on bacterial communities was analyzed for the three compartments of the digestive tract: the foregut (FG), midgut (MG), and hindgut (HG). However, due to poor results obtained for the FG (likely due to insufficient DNA quantity), the corresponding samples were excluded from the analyses. Therefore, our study focuses only on samples from the midgut and the hindgut. In all the results presented below, A corresponds to the fruit/vegetable waste diet, B to the potato scraps diet, and C to the forage diet. The sequences obtained in analysis of microbial ASVs from MG and HG samples were distributed in 997 ASVs (**Tables S2 & S3**), representing 12 phyla, 17 classes, 47 orders, 84 families, and 161 genera (**Tables S4 & S5**). The number of ASVs, bacterial genera identified, and alpha diversity indices, i.e., the Shannon and Simpson index values, are presented in **Figure S1**. Alpha diversity metrics indicate that the hindgut harbors more balanced microbial communities, while the midgut exhibits greater taxonomic diversity but is more sensitive to diet. In terms of species richness (N_ASV and N_Genus), MG_C clearly dominates, suggesting that the diet C (forage) strongly promotes species richness in the midgut. MG_A has the lowest indices, particularly very low Shannon and Simpson indices (∼2.5 and ∼0.80), indicating a community dominated by a small number of taxa. Overall, diet has a greater influence on the structure of the midgut microbiota (where active digestion and physicochemical conditions vary depending on the substrates ingested), while the hindgut maintains a more robust diversity (likely due to a more anaerobic and stable environment promoting microbial fermentation).

Beta-diversity analyses, as highlighted by generalized principal coordinate analysis (PCoA), underscore the influence of diet and gut compartment at the levels of ASV, phyla, and families. They confirm the reliability of the methodology used, with consistent trends across the four biological replicates per condition (compact clusters). The MG and HG groups occupy distinct areas without any overlap, indicating that the gut compartment is the primary determinant of gut microbiota composition, far beyond the effect of diet (**Figures S2-4**). The effect of diet on microbiota composition is secondary but real, since, within each compartment, the three carbohydrate-rich diets (A, B, C) form distinct and separate scatter plots at all taxonomic levels (ASV, family, and phylum) (**Figures S2-4**). It should be noted that, for the midgut, the group corresponding to condition MG_B (100% potato scraps waste) tends to diverge significantly from the MG_A and MG_C conditions. In summary, beta-diversity analyses confirm that the midgut and hindgut harbor structurally distinct bacterial communities.

### 2. A highly plastic microbiota in the midgut and a more stable one in the hindgut

The main bacterial phyla detected in all conditions are: Proteobacteria, Firmicutes, Bacteroidota, and Actinobacteriota. The least common phyla include Campylobacterota, Desulfobacterota, Fusobacteriota, Planctomycetota, Patescibacteria, Verrucomicrobiota, WPS-2, and Cyanobacteria (**Figure S5)**. What stands out most from the examination of the bacterial community analyses is that sequence abundance is particularly high for the MG_A condition (*i.e.,* 100% fruit/vegetable waste diet at the midgut level) compared to the other conditions (**Figures S5-7**). The condition MG_A is overwhelmingly dominated by *Actinomyces* (Actinobacteriota), a genus also presents in the other conditions, but at much lower abundance.

The analyses show that most of the identified bacterial phyla are present regardless of the conditions, but that their relative abundance can vary considerably from one condition to another (**Figure 1**). For instance, the phylum Actinobacteriota is well represented in all treatment groups, but its relative abundance is significantly higher in the MG_A condition (at the expense of the phylum Firmicutes compared to the MG_B and MG_C conditions), *i.e*., in the midgut of BSFL fed a diet consisting of 100% fruit/vegetable waste. The phylum Bacteroidota is present under all conditions, but its relative abundance is highest in the hindgut, regardless of diet. Overall, analyses conducted at the phylum level tend to confirm that the microbiota of the hindgut is more stable than that of the midgut, regardless of diet.

**Figure 1.**
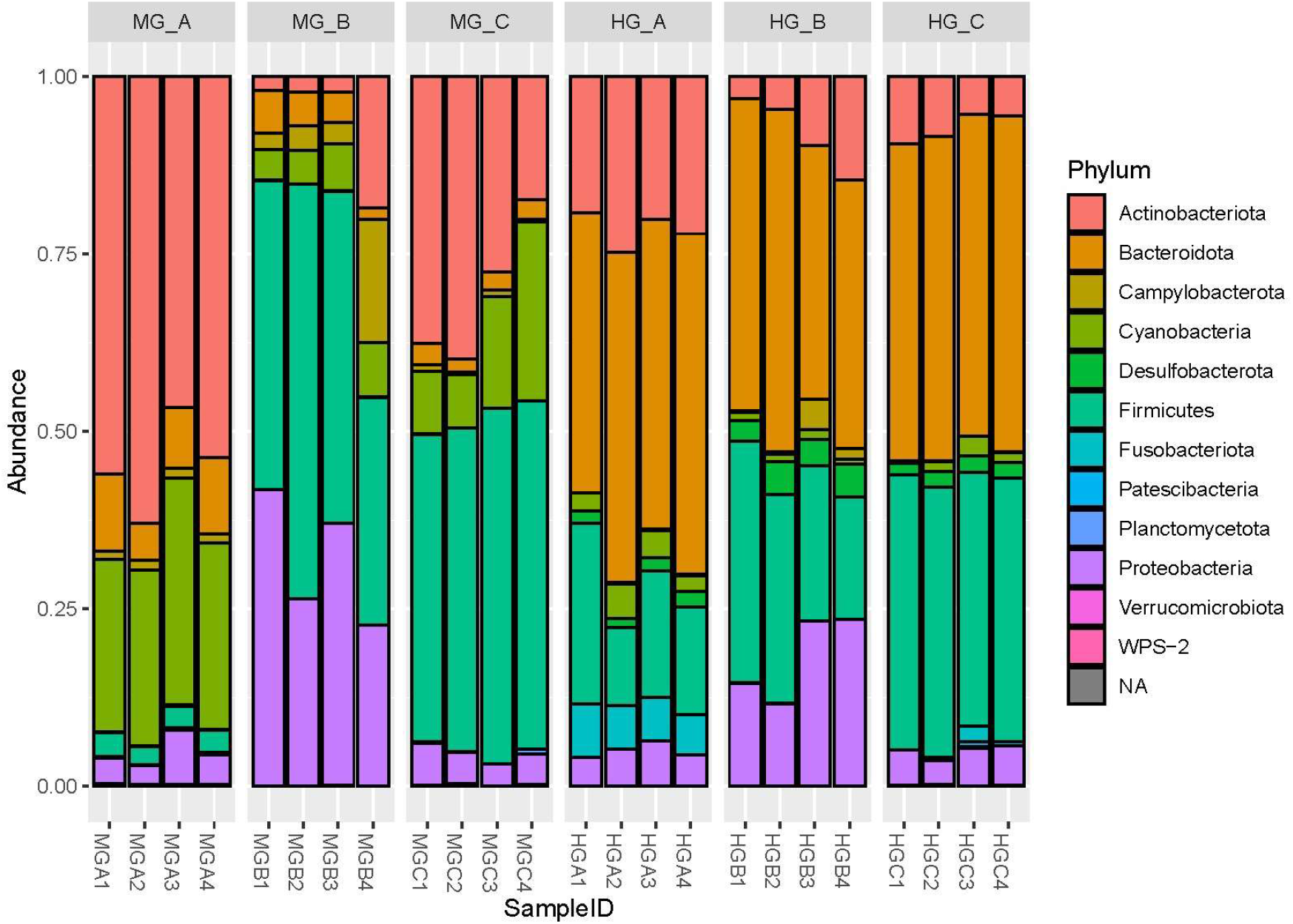
Relative abundance of bacterial phyla in the midgut (MG) and hindgut (HG) of BSFL fed on three distinct carbohydrate-rich diets: 100% fruit/vegetable waste (A), 100% potato scraps (B), and 100% forage (C). Metabarcoding analyses were performed on four biological replicates (1-4).

Given the large number of bacterial genera identified (**Figure S8**), we focused on the 10 most abundant for greater clarity (**Figure 2**). These are *Actinomyces*, *Bacillus*, *Bacteroides*, *Campylobacter*, *Clostridium*, *Desulfovibrio*, *Dysgonomonas*, *Fusobacterium*, *Morganella*, and *Paenibacillus* were identified as the 10 genera with the highest relative abundance. It is important to note that a significant portion of the bacterial diversity could not be identified (“undetermined”) and that a large portion of the identified bacterial diversity comprises a range of genera not included in the top 10 (“other”). Overall, the relative abundance of the top 10 bacterial genera varies across conditions, with marked trends. For instance, compared to the microbiota of the hindgut, that of the midgut shows the most pronounced response to substrate composition, with communities differing markedly between dietary treatments. The midgut microbiota of BSFL reared on fruit and vegetable waste (MG_A) is overwhelmingly dominated by *Actonimyces* while the MG_B microbiota (100% potato scraps waste) is featured by a high proportion of unassigned sequences. The midgut of BSFL reared on forage (MG_C) is featured by the prominent presence of *Paenibacillus* spp. alongside *Actinomyces* spp. and *Bacillus* spp. In sharp contrast to the substrate-sensitive midgut, the hindgut microbiota shows a markedly convergent and stable composition across all dietary treatments, dominated primarily by the genus *Dysgonomonas* (approximately 50% abundance). The results show similar patterns across biological replicates, confirming the robustness of our methodology. Overall, high-carbohydrate diets have a significant influence on the diversity and relative abundance of the bacteria that make up the BSFL microbiota. However, this influence is more pronounced for the midgut microbiota than for the hindgut microbiota, indicating once again that the hindgut harbors a more stable microbiota than the midgut.

**Figure 2.**
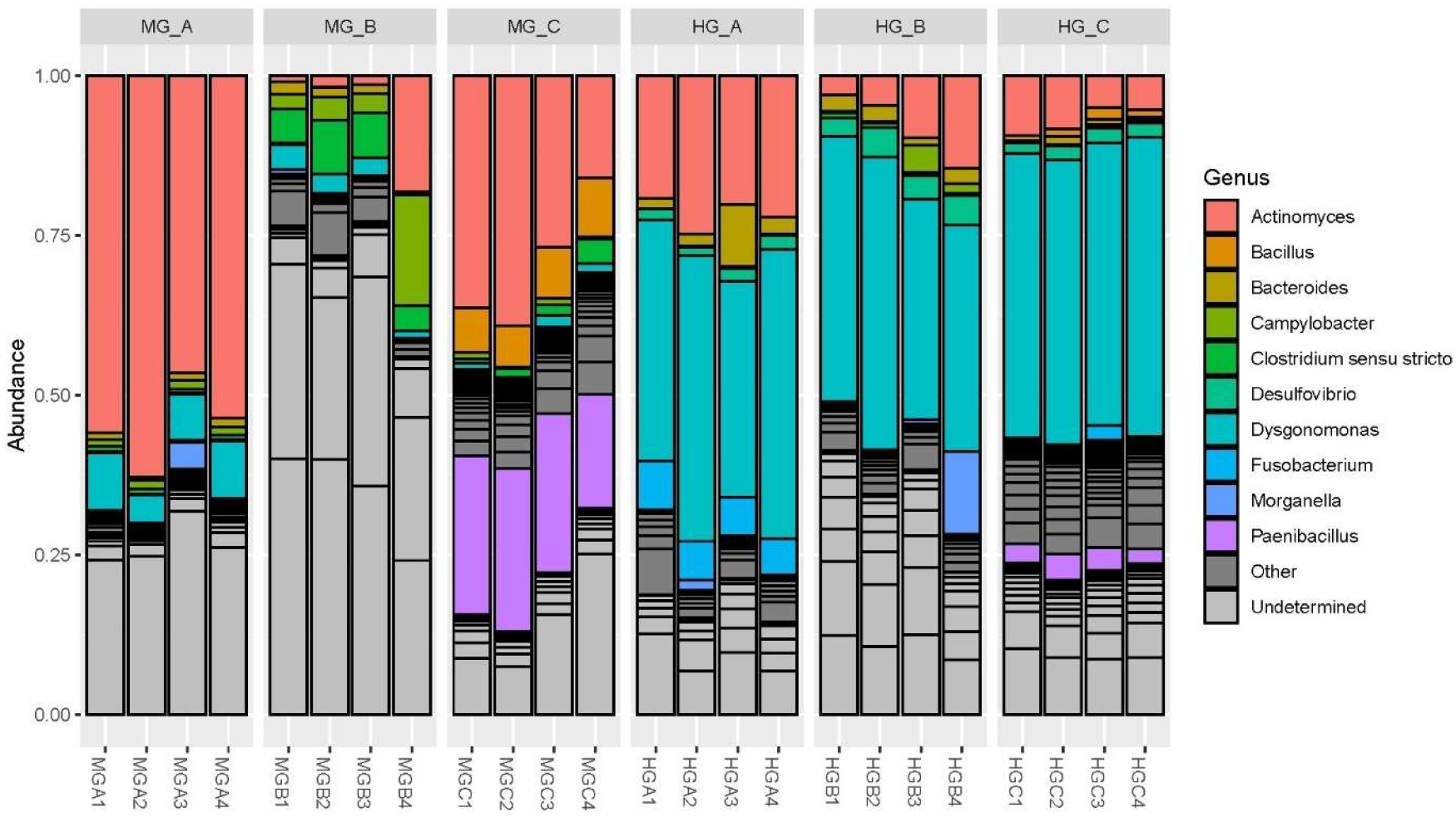
Relative abundance of the 10 most abundant bacterial genera in the midgut (MG) and hindgut (HG) of BSFL fed on three distinct carbohydrate-rich diets: 100% fruit/vegetable waste (A), 100% potato scraps (B), and 100% forage (C). Metabarcoding analyses were performed on four biological replicates (1-4).

We conducted additional analyses considering all identified genera to pinpoint those that were significantly more abundant under one treatment condition compared to the others (one-*versus*-all approach) (**Table S5 & Figure 3**). While many of these taxa do not rank among the top 10 in terms of relative abundance, they nonetheless remain significant potential contributors to the digestion of organic matter and serve as important indicators of the functions performed by the bacterial communities in each compartment of the BSFL digestive tract. The MG_A condition is the one with the highest number of significantly enriched genera, all on the right (positive CLR difference): *Comamonas*, *Pectobacterium*, *Leuconostoc*, *Lactococcus*, *Flavobacterium*, *Myroides*, *Ruminococcus*, etc. On the left (depleted in MG_A), there are anaerobic genera such as *Clostridium* and *Herbinix*. MG_A therefore has a very distinct microbial signature, dominated by aerobic/aerotolerant bacteria. The MG_B group includes numerous enriched genera, such as *Neoscardovia*, *Paucilactobacillus*, *Schleiferlactobacillus*, *Bifidobacterium*, *Lacticaseibacillus*, and others. There is a strong presence of Lactobacillales and lactic acid bacteria, suggesting that diet B (potato scraps) promotes the growth of bacteria capable of converting sugars into lactic acid through fermentation in the midgut. The MG_C condition exhibits the highest number of significant enriched genera, including *Devosia*, *Aminobacter*, *Rhodococcus*, *Paenibacillus*, *Mycobacterium*, and many others. Diet C therefore induces the most complex and diverse midgut community, which is consistent with the alpha diversity results. The HG_A condition contains few significant genera overall. The enriched genera include *Fusobacterium*, *Pseudomonas*, *Lachnoclostridium*, and *Anaerotruncus*. With the exception of *Pseudomonas*, these are strict anaerobic bacteria typical of the hindgut. The HG_B condition exhibits one significantly enriched genus: *Orbus*, a little-known genus. The HG_C condition is characterized by an enrichment of the Christensenellaceae R-7 group and *Anaerovorax* (strictly anaerobic bacteria), as well as several genera associated with fermentation (*Tyzzerella*, *Monoglobus*, *Ruminiclostridium*), indicating increased fermentative activity in the hindgut under diet C.

**Figure 3.**
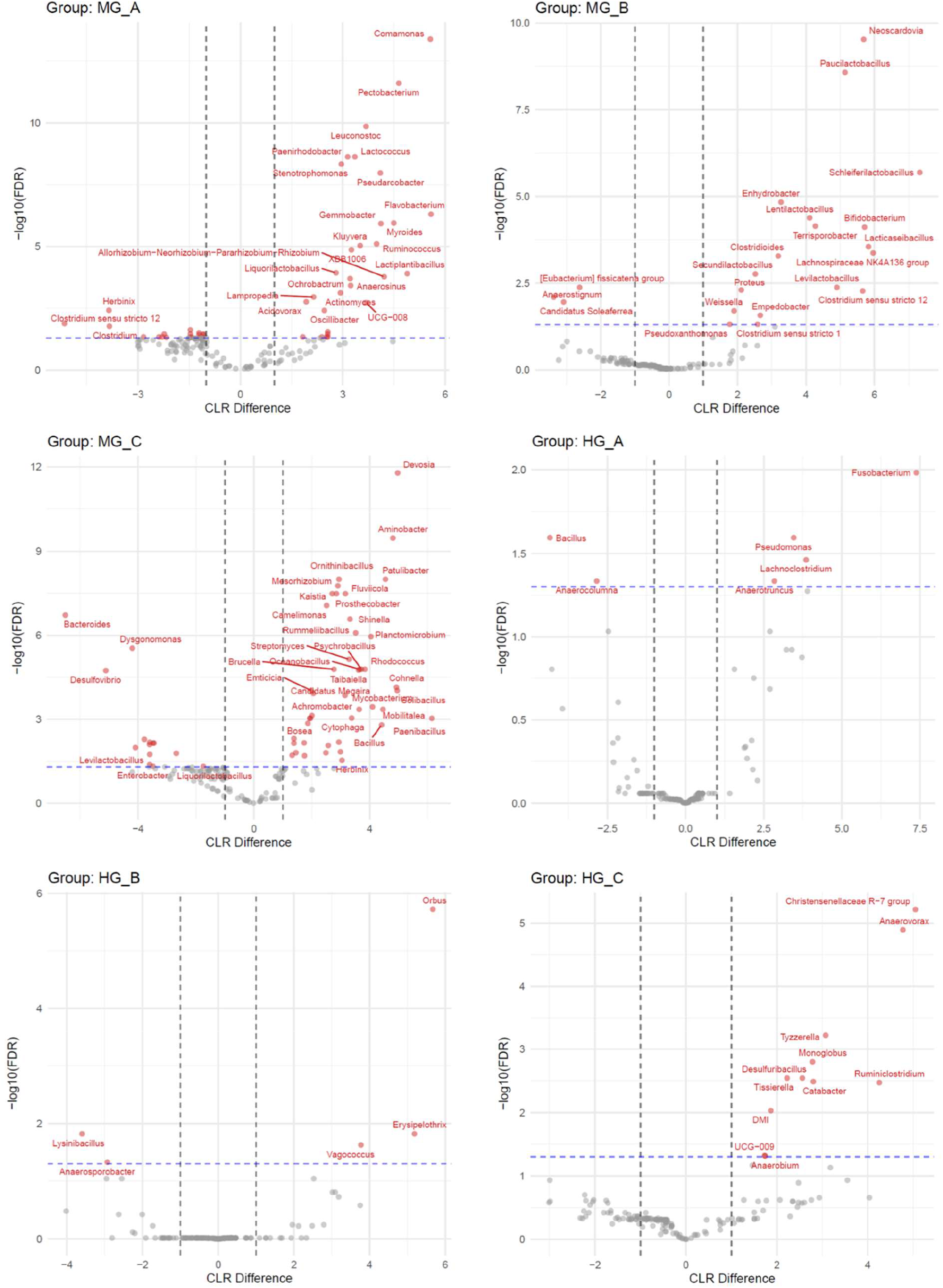
Differential abundance analysis of bacterial genera in the midgut (MG) and hindgut (HG) of BSFL reared on three different diets (A, B and C). Volcano plots represent, for each group (MG_A, MG_B, MG_C, HG_A, HG_B, HG_C), the difference in bacterial abundance expressed as centered log-ratio (CLR difference, x-axis) against statistical significance (−log10 FDR, y-axis). Each dot corresponds to a bacterial genus. Red dots indicate significantly differential genera (FDR < 0.05, horizontal blue dashed line) with an effect size exceeding the defined threshold (vertical dashed lines). Positive CLR values indicate enrichment in the focal group, whereas negative values indicate relative depletion. The most significant genera are annotated. n = 4 biological replicates per condition.

### 3. Prediction of the functional profiles of microbial communities

To further characterize the functional potential of the BSFL gut microbiota, metabolic pathways were inferred from 16S rRNA gene data using PICRUSt2 (**Figures 4 & S9**). Factorial analyses of CLR-transformed functional-category abundances revealed significant effects of diet on 10 of the 11 categories examined, with plant cell wall/lignocellulose degradation being the only exception, and significant effects of gut compartment on all categories except aromatic/phenolic compound degradation (**Table S1**). Significant diet x compartment interactions were detected for aromatic/phenolic compound degradation, energy metabolism/respiration, fatty acid biosynthesis/lipid metabolism, plant cell wall/lignocellulose degradation, starch/glycogen metabolism, and vitamin/cofactor biosynthesis (FDR-adjusted p < 0.03; **Table S6**). Within the midgut, diet C was associated with a higher predicted relative representation of amino acid and cell wall/envelope biosynthesis, energy metabolism, fermentation, simple-sugar degradation, starch/glycogen metabolism, and vitamin/cofactor biosynthesis than diets A and/or B, whereas amino acid degradation/nitrogen recycling was lower under diet C than under diets A and B (BH-adjusted p < 0.001) (**Table S6**). Diet B produced a distinct midgut profile characterized by a markedly higher relative representation of fatty acid biosynthesis and lipid metabolism than diets A and C (both BH-adjusted p < 0.001). In the hindgut, diets A and C showed similar profiles for several functional categories, whereas diet B was characterized by a higher predicted relative representation of aromatic/phenolic compound degradation than both other diets (both BH-adjusted p< 0.001) and lower fermentation, simple-sugar degradation, and starch/glycogen metabolism (BH-adjusted p < 0.005). Plant cell wall/lignocellulose degradation was highest in the hindgut under diet C, exceeding that observed under diets A and B (BH-adjusted p = 0.030 and 0.005, respectively). Overall, the midgut showed a greater relative representation of several biosynthetic, fermentative, and carbohydrate-related functions, whereas amino acid degradation/nitrogen recycling and plant cell wall/lignocellulose degradation were more strongly represented in the hindgut (**Table S7**).

**Figure 4.**
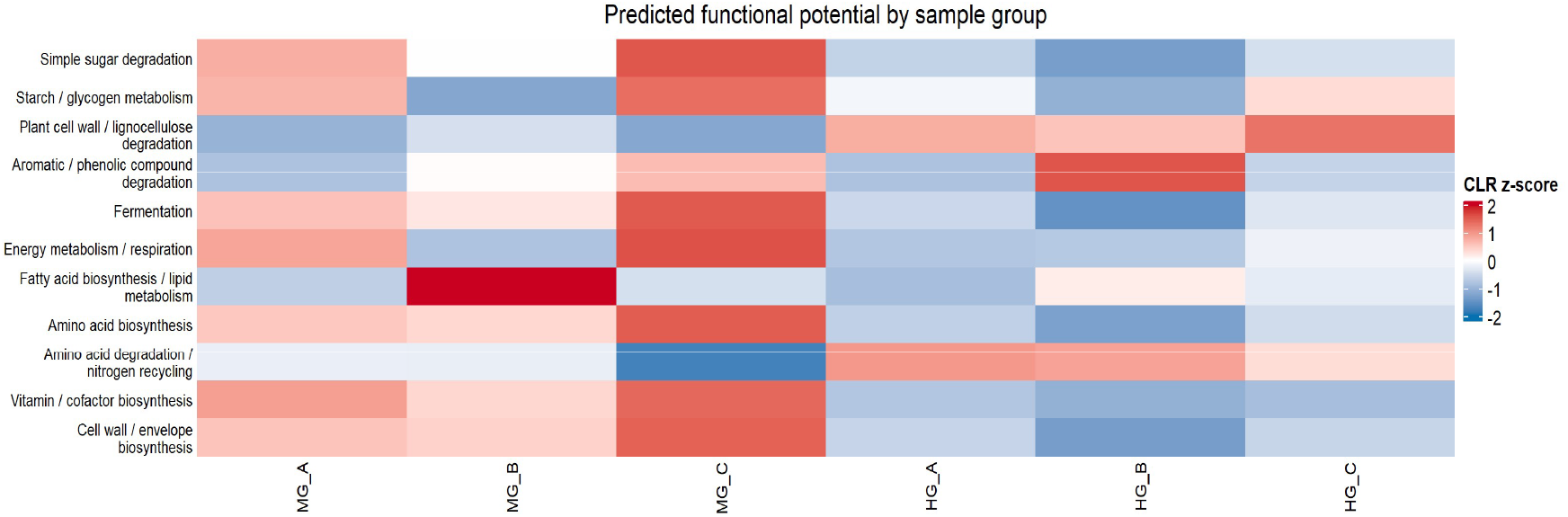
Predicted functional potential of the gut microbiota across diet–gut compartment groups. Functional profiles were inferred from 16S rRNA gene data using PICRUSt2, and predicted pathways were grouped into broad metabolic categories. Category abundances were centered log-ratio (CLR) transformed and standardized within each category as z-scores across sample groups. Red indicates a higher-than-average predicted relative abundance and blue a lower-than-average predicted relative abundance for a given functional category, whereas white indicates values close to the category mean. HG = Hind gut, MG = Mid gut, A corresponds to the fruit/vegetable waste diet, B to the potato scraps diet, and C to the forage diet.

## Discussion

BSFL are highly efficient bioconversion organisms capable of growing on a remarkably wide range of organic substrates. This polyphagous ability is primarily driven by a highly flexible gut microbiota, whose composition is modulated according to the chemical nature of the ingested substrate (20). Until now, the gut microbial community has been studied primarily at the whole-larva level, with comparatively few studies specific to the hindgut microbiota (22). The present results, based on 16S rRNA amplicon sequencing of the bacterial microbiota of the midgut (MG) and hindgut (HG) of BSFL reared on carbohydrate-rich diets exhibiting contrasting molecular compositions, provide evidence of a two-compartment microbial system in which the midgut acts as a substrate-responsive digestive compartment, whereas the hindgut harbors a less diverse and comparatively diet-stable bacterial community involved in the terminal processing of residual substrates and nutrient recycling.

Beta-diversity analyses confirm that the gut compartment is the primary determinant of microbial community composition, with the MG and HG samples forming completely distinct clusters at all taxonomic levels (ASV, family, phylum). This structural separation is consistent with the recent findings demonstrating that bacterial communities vary considerably between gut compartments, regardless of diet (22–24). However, these studies focused on the anatomical subdivisions of the midgut (anterior *versus* middle *versus* posterior) and did not address the hindgut, whereas our strategy was to consider the midgut as a whole for a comparative analysis with the hindgut. Our results highlight clear and distinct profiles in bacterial composition between the midgut and the hindgut, which are separated by the insertion of Malpighian tubules - a distinct anaerobic environment specialized in fermentative metabolism and nutrient recovery (22). The following sections examine the functional diversity of the bacterial microbiota in these two gut compartments, in light of the nature of the carbohydrate-rich diet.

An apparently unexpected result of the PICRUSt2 functional inference was the consistently higher relative representation of fermentation-related pathways in the midgut than in the hindgut. This pattern should be interpreted with caution, because the analysis does not measure direct pathway expression nor provides a mechanistic explanation. In addition, CLR-transformed values are relative: the lower representation of fermentation-related pathways in the hindgut may partly reflect the greater relative contribution of other predicted functions, including amino acid degradation, nitrogen recycling, and lignocellulose processing. Moreover, the BSFL midgut is an exceptionally long and heterogeneous organ. Its anterior region is an important site of polysaccharide hydrolysis, whereas bacterial abundance increases toward the posterior midgut. Because the entire midgut was analyzed as a single compartment, the observed functional profile may integrate the production of readily fermentable carbohydrates in upstream regions with the functional potential of the denser bacterial community occurring further downstream (21).

Our results show that the midgut microbiota exhibits the most pronounced response to substrate composition, with bacterial communities displaying significant differences across different diets; this finding supports the observation that dietary composition shapes the gut bacterial communities of BSFL (18, 29). The midgut of BSFL reared on the 100% fruit/vegetable waste diet (rich in carbohydrates, soluble fiber, and water; low in protein and fat) - that is, substrate A/condition MG_A - is overwhelmingly dominated by *Actinomyces* (Actinobacteriota), which account for more than 50% of the relative abundance across the four biological replicates. The genus was identified as the most common genus across all conditions, but it is by far the most dominant in the MG_A condition. The genus *Actinomyces* has been identified as a core member of the BSFL gut microbiota in multiple independent studies (30– 33). The exact role played by *Actinomyces* spp. in the digestion of organic matter by BSFL has yet to be fully elucidated. Members of this genus are Gram-positive, anaerobic or facultatively anaerobic bacteria that often exhibit a fermentative carbohydrate metabolism and are capable of breaking down sucrose into glucose and fructose (34). Consequently, its predominance in the midgut of BSFL fed exclusively on fruit and vegetable waste - the diet richest in sucrose, glucose, and fructose - is biologically consistent. Furthermore, it is well known that the order Actinomycetales includes versatile degraders of plant polysaccharides, including pectin and hemicellulose (35–37), which are abundant in fresh fruit and vegetable waste and produce short-chain organic acids (*e.g.,* acetate, succinate) that may contribute to the acidification of the midgut lumen (38). The second most abundant genus identified in MG_A is *Dysgonomonas* (accounting for approximately 10–15% of the relative abundance across the four biological replicates). Given the facultative anaerobic lifestyle of members of this genus, one would expect to find *Dysgonomonas* spp. in the hindgut rather than the midgut. Nevertheless, the presence of this genus in the midgut of BSFL has been previously confirmed; it may contribute to the degradation of complex polysaccharides and play a role in the degradation of lignocellulose (29, 39–41). Our study also identified numerous taxa with a low relative abundance that are specifically associated with MG_A. These include, among others, *Comamonas*, *Pectobacterum*, *Leuconostoc*, *Lactococcus* and *Ruminococcus*. The potential role of *Comamonas* spp. in the BSFL is difficult to pinpoint, but it is known that members of this genus are capable of metabolizing complex compounds such as lignin (42, 43). *Pectobacterium* spp. are necrotrophic, facultative anaerobic, Gram-negative plant pathogens, well known for producing a diversity of plant cell wall-degrading enzymes (PCWDE), such as pectinases, cellulases and proteases that break down plant tissues (44, 45). They tend to efficiently metabolize sugars, particularly sucrose and xylose. It is possible that, in the midgut of BSFL, *Pectobacterium* bacteria contribute to the breakdown of plant tissue during digestion. *Leuconostoc* spp. are heterofermentative lactic acid bacteria able to express a wide variety of carbohydrate-active enzymes (CAZymes), enabling them to colonize plant and dairy environments rich in complex carbohydrates (46, 47). *Lactococcus* spp. are known as homofermentative lactic acid bacteria (LAB), meaning they primarily metabolize hexoses (such as glucose, galactose, and lactose) into lactic acid as the main end product (48). *Ruminococcus* spp. include key gut bacteria that specialize in breaking down complex carbohydrates, such as resistant starch, cellulose, and hemicellulose (49, 50). The genus *Ruminococcus* is well documented in mammals (particularly ruminants), and its presence in the BSFL midgut fed a complex, carbohydrate-rich diet is therefore consistent (49–51). Overall, our results show that the bacterial microbiota associated with MG_A consists mainly of *Actinomyces* and *Dysgonomonas*, as well as a range of satellite bacteria primarily involved in carbohydrate metabolism. This taxonomic profile is consistent with the higher predicted relative representation of pathways related to sucrose degradation and carbohydrate fermentation under the MG_A condition (27).

The most striking feature of the MG_B microbiota is the high proportion of unassigned sequences (60-75% “Undetermined” at genus level across all replicates) compared with the MG_A and MG_C conditions. This condition exhibits the most marked differences in terms of bacterial composition and metabolic activity. Potato scraps contain high concentrations of resistant starch, but also potentially significant amounts of glycoalkaloids (*e.g.,* solanine, chaconine) in the scraps fraction, which may exert broad-spectrum antimicrobial activity (52). One hypothesis is that the selective pressure imposed by these antinutritional compounds over 13 days of continuous exposure likely eliminates susceptible taxa and enriches a community of resistant or detoxifying specialists that are poorly represented in current reference databases. We propose that the taxonomic novelty of the MG_B bacterial community represents a potentially significant reservoir of unexplored enzymatic and detoxification capabilities that warrant further characterization (*e.g.,* via shotgun metagenomics). Among the most abundant genera identified are *Clostridium sensu stricto* and *Campylobacter*. *Clostridium sensu stricto* spp. are strictly anaerobic, fermenting bacteria that exhibit a saccharolytic activity and utilize carbohydrates (glucose, xylose, etc.) for growth (53). *Campylobacter* spp. are microaerophilic, chemoorganotrophic bacteria with restricted, non-saccharolytic metabolism, meaning they do not use carbohydrates (like glucose) for energy. Instead, they rely on amino acids (serine, aspartate, glutamate) and organic acids (pyruvate, fumarate, acetate) as primary energy sources. It is interesting to note that the bacteria specifically associated with MG_B include a diversity of lactic acid bacteria (*e.g., Paucilactobacillus*, *Schleiferilactobacillus*, *Lentilactobacillus*, *Lacticaseibacillus*, and *Bifidobacterium*). This suggests that carbohydrates are fermented into lactic acid, which may lead to a decrease in the pH of the midgut. The high fatty acid content of substrate B (27), which is particularly conducive to the growth of these bacteria, likely explains their specific presence in the midgut of BSFL. Another important point to note is that BSFL raised on a 100% potato scraps diet tend to have significantly higher fatty acid content than those fed diets consisting of 100% fruit/vegetable waste and 100% forage (27). This implies that the microbiota specific to the MG_B condition plays a significant role in fatty acid metabolism, although the direction of causality and the actual contribution of bacterial metabolism cannot be determined from the present data. The absence of highly dominant and well-known genera makes it difficult to attribute metabolic functions to specific genera. However, metabolic activity oriented toward fatty acid biosynthesis is clearly confirmed by the PICRUSt2 functional analysis (*e.g.,* superpathway of fatty acid biosynthesis initiation, palmitoleate biosynthesis, oleate biosynthesis, stearate biosynthesis, palmitate biosynthesis and mycolate biosynthesis). In addition, substrate B is richer in fatty acids than substrates A and C, and this richness is reflected in the composition of the larvae (27). In light of these results, potato scraps waste constitutes an optimal substrate to customize BSFL for use in, for example, aquaculture, livestock feed and cosmetics (54–57).

The midgut of BSFL reared on forage (MG_C), which is rich in lignocellulose, is characterized by high alpha diversity (highest N_ASV and Shannon index in the entire dataset) and a prominent presence of *Paenibacillus* (Firmicutes) alongside *Actinomyces* and *Bacillus*. *Paenibacillus* spp. are among the most versatile lignocellulose-degrading bacteria known (58), encoding multiple carbohydrate-active enzymes (CAZymes) including cellulases, hemicellulases, xylanases, and pectinases (59–62). The enrichment of the genus in the midgut of forage-fed BSFL directly reflects the lignocellulosic nature of the substrate and represents a functional convergence with the rumen microbiome of ruminants, where *Paenibacillus* plays analogous roles in the degradation of plant cell walls (63, 64). In addition to these dominant genera, a multitude of bacterial genera specific to MG_C have been identified. Unfortunately, this diversity includes a significant proportion of bacteria described as being associated with soil and plants, but which are for the most part poorly documented metabolically (*e.g., Devosia*, *Aminobacter*, *Ornithinibacillus*, *Patulibacter*, *Mesorhizobium*, *Fluviicola*, *Kaistia*, *Rummeliibacillus*, etc.). It is therefore difficult to pinpoint the role of these genera in forage digestion. However, among this diversity, *Cellulosimicrobium* stands out from a functional perspective: it is a genus of Gram-positive bacteria belonging to the order Actinomycetales, primarily known for its highly efficient ability to break down plant material through its lignocellulolytic activity, and which has been identified as a gut symbiont of termites (65–67). Although MG_A and MG_C shared several qualitative features in the heatmap, the factorial analysis revealed a distinct predicted functional profile under MG_C. In particular, amino acid biosynthesis was more strongly represented under MG_C than under MG_A and MG_B, together with higher predicted representation of several energy-, fermentation-, carbohydrate-, and vitamin-related functions. Conversely, amino acid degradation and nitrogen recycling were less strongly represented under MG_C than under the other midgut conditions. Whether this predicted microbial biosynthetic potential contributes to the higher protein content previously observed in forage-fed larvae remains to be determined, but could explain what makes forage-fed black soldier flies have a higher protein content (27). Overall, our study highlights the pivotal role of *Paenibacillus* spp. in forage digestion, thanks to their ability to produce enzymes capable of breaking down the complex carbohydrates found in this substrate.

In sharp contrast to the substrate-sensitive microbiota of the midgut, that of the hindgut exhibits a distinctly stable and homogeneous composition, regardless of diet. The dominant taxon, consistently and overwhelmingly, is *Dysgonomonas*, which accounts for approximately 50% of the relative abundance in all conditions, regardless of whether the larvae were fed fruit/vegetable waste, potato scraps, or forage. This substrate-independence identifies *Dysgonomonas* unambiguously as the keystone taxon of the BSFL hindgut. The genus *Dysgonomonas* comprises facultatively anaerobic, fermentative bacteria of the family Dysgonomonadaceae (phylum Bacteroidota). *Dysgonomonas* spp. have been isolated from diverse environments, including human clinical samples, sea sand, microbial fuel cells, human gut and the hindgut of termites (68–74). The functional importance of the genus in the BSFL gut is among the best-documented in the literature, despite its relatively recent characterization. *Dysgonomonas* members have been identified as representatives of the BSFL core gut community, present in larval guts irrespective of the waste type used as a rearing substrate (18, 20, 75–78). *Dysgonomonas* spp. are primarily known to be associated with the hindgut of termites and likely play a key role in the digestion of complex lignocellulosic polysaccharides, as many species also possess a wide array of carbohydrate-active enzymes (CAZymes) (79– 81). Functionally, the presence of *Dysgonomonas* spp. in the BSFL gut is positively correlated with genes involved in carbohydrate, sulfate, and nitrogen metabolism (82). Furthermore, a metagenomic analysis of the BSFL gut traced the origin of a novel α-galactosidase gene enabling the hydrolysis of α-galactoses abundant in non-digestible plant oligosaccharides to a specific *Dysgonomonas* strain (83). More recently, *Dysgonomonas* spp. were identified as strongly correlated with protein digestion and absorption in BSFL reared on protein-rich artificial diets (76). On the basis of the available literature and the present data, *Dysgonomonas* spp. can be assigned a tripartite functional role in the hindgut of BSFL. The first role is secondary polysaccharide degradation: thanks to their α-galactosidase and broader CAZyme repertoire, *Dysgonomonas* bacteria could process residual plant-derived polysaccharides that escaped midgut digestion, thereby maximizing the extraction of fermentable sugars in the terminal gut compartment. This function is consistent with the enrichment of *Dysgonomonas* and hemicellulolytic enzyme families (GH51, GH43_16) in BSFL fed lignocellulosic diets (39). Another role involves nitrogen and sulfate metabolism: the positive correlation of *Dysgonomonas* spp. with genes involved in nitrogen and sulfur metabolism (82) suggests a role in ammonium production via deamination of residual amino acids and in sulfur redox cycling in the hindgut environment, which contribute to the broader nutrient recycling capacity of the larva. Finally, a third role could be protein digestion and absorption: its direct implication in protease-mediated protein breakdown and amino acid absorption (76) makes *Dysgonomonas* a multifunctional digestive partner, operating beyond carbohydrate metabolism to enhance overall nitrogen assimilation efficiency. The substrate-independence of *Dysgonomonas* dominance across conditions A, B, and C, spanning chemically very distinct substrates, is the most compelling evidence in the present dataset for its status as a true core taxon whose ecological niche is defined by gut physiology rather than dietary input. To our knowledge, although *Dysgonomonas* has previously been described as a pervasive bacterial associate in BSFL, this is the first time a clear link has been established between the presence of *Dysgonomonas* and the functioning of the hindgut. The PICRUSt2 functional analysis suggests metabolic activity geared towards the degradation of lignocellulose and amino acids and oriented toward nitrogen recycling - roles likely performed primarily by *Dysgonomonas* spp.

Unlike the midgut, the hindgut has extremely low bacterial diversity, regardless of the conditions. *Fusobacterium*, a Gram-negative anaerobic genus, is clearly specific to the HG_A condition, but its function remains unknown. One hypothesis is that it is capable of fermenting both amino acids and glucose (84). The genus *Orbus*, discovered relatively recently, is specific to the HG_B condition and comprises Gram-negative, facultative anaerobic bacteria frequently found in the insect gut, but with no clear indication of their presumed role (85–89). The Christensenellaceae R-7 group is a genus of bacteria belonging to the family Christensenellaceae (phylum Firmicutes) that has been identified as a key component of the gut microbiota of various insects, particularly those that feed on wood or decaying plant matter, such as the larvae of Cerambycidae, Scolytidae, and Lucanidae (90–92). These bacteria are generally associated with cellulose metabolism and the degradation of plant material, thereby helping their hosts extract nutrients from a fibrous diet. The presence of this genus in the hindgut of forage-feeding BSFL is therefore consistent; it may act synergistically with the genus *Dysgonomonas* in the digestion of lignocellulose, thereby enabling the insects to utilize nutrient-poor plant material.

Extensive research has been conducted on the composition of the gut microbiota of BSFL (20). However, we are only just beginning to understand how it is distributed across the different compartments of the digestive tract. And while the midgut has been the subject of a few recent studies (17, 21–24), the hindgut remains a black box. By focusing on three diets rich in carbohydrates but with contrasting molecular compositions, our analyses revealed that the functional architecture of the gut microbiota of BSFL can be summarized in a revised two-compartment model: the midgut acts as a chemically responsive adaptive chamber, in which substrate-specific microbial guilds are recruited in response to the biochemical nature of the ingested material. Its community is plastic and substrate-dependent. The hindgut, by contrast, maintains a less diverse and comparatively diet-stable community dominated by *Dysgonomonas* spp. regardless of upstream dietary composition. While the midgut handles the initial breakdown, and appears to combine substrate-responsive digestion with a relatively high predicted representation of fermentative, biosynthetic, and carbohydrate-related functions, the hindgut tends to process the remaining undigested material through anaerobic fermentation. It is more strongly characterized by predicted functions associated with the processing of residual lignocellulosic and nitrogenous compounds. This revised model therefore differs from the conventional view of the hindgut as the principal fermentation chamber. Nevertheless, because PICRUSt2 estimates relative genomic potential rather than metabolic activity, the spatial distribution of fermentation within the BSFL gut will need to be validated using compartment-specific metabolomic, metatranscriptomic, or direct biochemical measurements.

The more complex the substrate, the richer and more diverse the overall bacterial community tends to be. However, it remains difficult to precisely determine the role played by each bacterial partner in this process. Few genomes of associated gut bacteria are currently available (including those of dominant taxa). Yet this is an essential step toward moving beyond functional hypotheses regarding their role in insect digestion and better understanding the larvae’s remarkable ability to feed on a wide variety of organic materials. Culture-based approaches are crucial for accurately capturing the functional diversity of the bacterial microbiota of BSFL and assigning a metabolic role to associated bacteria based on their sequenced and annotated genomes. This is also an important step toward harnessing this insect’s highly adaptable microbiome for industrial purposes, as well as toward understanding how an insect is able to digest such a wide variety of organic substrates. The present study represents a step forward in this direction.

## Material and methods

### 1. Preparation of the carbohydrate-rich diets

For the experiments, the larvae were reared on three different carbohydrate-rich diets prepared as described previously (27): 1) 100% of fruit/vegetable waste, 2) 100% of potato scraps, and 3) 100% of forage. The fruit/vegetable waste and potato scraps were collected from the university canteen (Louvain-la-Neuve, Belgium). The fruit/vegetable waste included zucchini, carrots, celery, onions, and melons. The potato scraps included rotten potatoes, potato peelings, rotten sweet potatoes, and sweet potato peelings. Once collected, the waste was weighed and stored at -20°C until use. The forage consisted of a mixture of two plants: clover (Fabaceae) and ryegrass (Poaceae), which were grown and stored at the Alphonse de Marbaix experimental farm (UCLouvain, Belgium). The chemical profile of the three diets (crude protein content, amino acid profile, ash, mineral composition, sugar content, and fatty acid profile) was comprehensively determined in Eke *et al.* 2025 (27). Before use, the frozen substrates were removed from the freezer, thawed at room temperature for five hours, and then drained to remove excess water.

### 2. Larvae rearing conditions

The eggs were collected from a black soldier fly (BSF) stock colony established in May 2022 at the Alphonse de Marbaix experimental farm. The colony was maintained under controlled conditions of 28 ± 2 °C and 60 ± 5% relative humidity, as described by (93). The collected eggs were placed in small transparent plastic pots containing a mixture of 50 g of corn flour and 100 ml of water. The pots were covered with a fine mesh cloth and secured with rubber bands to prevent the young larvae from escaping. The entire setup was then placed in an incubator (Memmert) set to 28 ± 2 °C and 70 ± 5% relative humidity to promote proper incubation and hatching of the eggs. Daily observations were made to monitor hatching. After four days, the newly hatched larvae emerged. These neonates were then fed for five days using the cornmeal provided for hatching. On the fifth day after hatching, the larvae were separated from the corn-meal residue by sieving. A total of 3,600 larvae were counted manually and distributed among twelve separate rearing boxes with perforated lid (11L - IPL 4412, 40×27×5.5 cm, Iris Europe). Each box, containing 1 kg of feeding substrate, was inoculated with 300 five-day-old larvae, for a total of 4 replicates per diet.

### 3. Sample collection

The larvae were reared for 13 days, after which five larvae were randomly selected for each replicate using a forceps (tip width: 2.5 mm; length: 12 cm). The larvae were rinsed with tap water to remove substrate residues, gently dried with paper towels, and then inactivated by freezing at -20 °C for 15 minutes. They were then dissected to isolate the gut sections for metabarcoding analysis.

### 4. Gut dissection and DNA extraction

For the dissection of the digestive tract, after inactivation, the larvae were rinsed successively with clean water, distilled water, and 70% ethanol, in accordance with the procedure described by Wynants *et al*. (2019) (94). Each individual was then placed on a sterile Petri dish in a horizontal laminar flow hood. The digestive tract was dissected in sterile phosphate-buffered saline (PBS) using sterilized forceps and scalpels under a microscope to isolate the digestive tract and clearly identify its distinct anatomical parts. To minimize the risk of external contamination, a new sterile Petri dish was used for the dissection of each larva’s gut. Once the entire digestive tract had been removed, it was transferred to another sterile Petri dish containing PBS using a pair of sterile forceps. The tract was then carefully dissected into three main sections - the foregut, the midgut, and the hindgut - using sterile forceps. Any breakage or contact of the digestive tract with the rest of the larva’s body or the work surface during dissection resulted in the specimen being discarded. Additionally, any samples that came into contact with each other during the separation process were discarded. All dissection tools were sterilized between uses by passing them through a Bunsen burner and cleaning them with 70% ethanol, in accordance with the protocol described by Lanan *et al*. (2016) [31]. For the subsequent metabarcoding analyses, five foreguts (FG), two midguts (MG) and two hindguts (HG) were collected to generate one replicate. In total, four replicates per gut region and diet were prepared. The dissected digestive tract segments were placed in separate Eppendorf tubes and stored at -80°C until use. DNA extraction was then performed using the DNeasy PowerSoil Pro kit (Qiagen) according to the manufacturer’s protocol.

### 5. Illumina metabarcoding of gut bacterial communities

The purity of the samples was validated using Nanodrop, and the integrity of the genomic DNA was validated using electrophoresis on a 1% agarose gel. The DNA concentration of the samples was measured using a Qubit fluorometer (Invitrogen, Gent, Belgium) with a Qubit dsDNA HS Assay kit (DNA concentration varied between 5 and 25 ng/µl). AMPure XP beads (Beckman Coulter, Indianapolis, IN) were used to remove contaminants from the 16S DNA. Samples were then diluted using HyClone™ water in equal concentrations (5 ng/µl) and stored at -20°C until high-throughput sequencing (HTS).

Sequencing libraries were prepared according to the Illumina 16S metagenomic sequencing library preparation protocol (San Diego, CA) at the Center of Applied Molecular Technologies (CTMA, UCLouvain, Brussels, Belgium). In brief, the V3-V4 variable region of the 16S bacterial rRNA gene was amplified using a two-stage PCR protocol. The first-stage PCR (PCR1) amplification uses universal primers of the interest region with overhang adapters attached (F: 5′TCG TCG GCA GCG TCA GAT GTG TAT AAG AGA CAG CCT ACG GGN GGC WGC AG and R: 5′ GTC TCG TGG GCT CGG AGA TGT GTA TAA GAG ACA GGA CTA CHV GGG TAT CTA ATCC) to amplify a 483-bp portion of the V3-V4 region. This step uses a kit 2x KAPA HiFi HotStart ReadyMix in a mixture total volume of 25 μl per sample. The PCR1 was run under the following conditions: 3 min denaturation at 95°C; 25 cycles of denaturation (30 sec at 95°C), annealing (30 sec at 55°C) and extension (30 sec at 72°C); a final extension at 72°C for 5 min. The PCR1 product was cleaned with AMPure XP beads in a 0,8X ratio (Beckman Coulter, Indianapolis, IN) to purify the 16S V3-V4 amplicon from free primers and primer dimer species. The second-stage PCR (PCR2) was done with 5 μl PCR1 purified to attach dual indices and Illumina sequencing adapters using the Nextera XT v2 Index Kit D. The PCR2 was performed under the same conditions as PCR1 with only eight PCR cycles of denaturation/annealing/extension. A clean-up of the PCR2 products with AMPure XP beads (Beckman Coulter, Indianapolis, IN) was performed before quantification on a Qubit fluorometer (Invitrogen, Gent, Belgium). All library sizes were then evaluated by capillary electrophoresis on an Agilent Technologies 2100 Bioanalyzer using an Agilent DNA 1000 kit (Agilent Technologies). After quantification and size determination, all libraries were equimolarly pooled to a final concentration of 4 nM. After NaOH denaturation and dilution, a 9 pM pooled library was loaded on MiSeq reagent kits v3 (600 cycles) including Illumina 15% PhiX control spike-in. Blanks consisting of sterile water instead of genetic material were performed to validate each laboratory step and to identify possible contamination (extraction, amplification and sequencing).

### 6. Bioinformatics analyses

All NGS reads were processed using R 4.2.1 and the dada2 Bioconductor package (96). Sequences were trimmed and filtered to tolerate a maximum of 2 expected errors per paired ends read. Amplicon sequence variants (ASV) were inferred using the high-resolution DADA2 method, which distinguishes sequencing errors from real biological variation. Chimeras and low abundance ASV making up <0.001% of reads were subsequently removed from the data set. Data from blank samples were analyzed to control for potential contamination in DNA extraction kits, as previously recommended (97). Taxonomy was assigned with a naive Bayesian classifier implemented in the DADA2 package and using the SILVA v.138.1 training set.

To assess α-diversity within the different microbial communities, richness, Shannon, and Simpson indices were calculated using the *phyloseq* Bioconductor package (98) after rarefying samples to a uniform sequencing depth of 50.000 reads.

Amplicon sequence variants (ASVs) were agglomerated at the phylum and genus levels using the *phyloseq* package, and relative abundances across different diets and compartments were visualized using barplots. To identify differentially abundant taxa, genus-level agglomerated data were subjected to a centered log-ratio (CLR) transformation. These scaled data were then analyzed using two-way ANOVA models to assess the main effects of diet and compartment. Concurrently, a one-versus-all approach was implemented via one-way ANOVA models to identify specific genera uniquely associated with distinct diet and compartment combinations. P-values were adjusted for multiple testing using the Benjamini-Hochberg False Discovery Rate (FDR) procedure.

To investigate differences in community composition (β-diversity), Bray-Curtis distances were calculated with the phyloseq R package and plotted using principal coordinates analysis.

PICRUSt2 (v2.1.4) was used to infer the functional potential of the gut microbial communities from amplicon sequence variants (ASVs) (99). Predicted pathway abundances were aggregated into broad functional categories by manually assigning pathway descriptions using a keyword-based classification of the pathway annotations (**Table S1**). Within each sample, pathway abundances were summed by functional category. Because the resulting data were compositional, a pseudocount of 1 was added before applying a centered log-ratio (CLR) transformation. For each functional category, CLR-transformed abundances were analyzed using factorial models including diet, gut compartment, and their interaction as explanatory variables. The significance of each term was assessed using Type II ANOVA, and P-values were adjusted for multiple testing using the Benjamini–Hochberg false discovery rate procedure. When significant effects or interactions were detected, pairwise comparisons among estimated marginal means were performed with Benjamini–Hochberg adjustment. For visualization, category-level CLR values were standardized within each functional category as z-scores. Heatmaps were generated in R using the *ComplexHeatmap* package, with rows representing functional categories and columns representing diet–compartment groups arranged in a predefined order. An additional heatmap displayed midgut and hindgut samples in separate panels. Functional categories were arranged in a biologically meaningful sequence, without hierarchical clustering of rows or columns. As a complementary exploratory analysis, one-versus-rest comparisons were used to identify functional categories showing particularly distinctive relative abundances within individual diet–compartment combinations. These comparisons were not used as the primary basis for statistical inference.

## Declarations

### Ethics approval and consent to participate

Not applicable.

### Consent for publication

Not applicable.

### Availability of data and materials

The NGS datasets generated during this study are available in the European Nucleotide Archive (ENA) repository under accession code PRJEB121040.

### Authors’ contributions

M.E, TH and F.R. designed this project. TH has secured funding for the research. M.E. and F.R. performed the study, analyzed the data, and wrote the paper. B.B. performed the sequencing. M.M. and J.A. generated the metabarcoding data and contributed to data analysis. L.S.N.T, K.T., M.M., J.A., B.B., J.L.G. and T.H. edited and revised the paper. All authors read and approved the final manuscript.

## Acknowledgements

This paper is publication BRC 443 of the Biodiversity Research Centre (Université catholique de Louvain).

## Supplemental material file: legends for supplementary Figures

**Figure S1.**
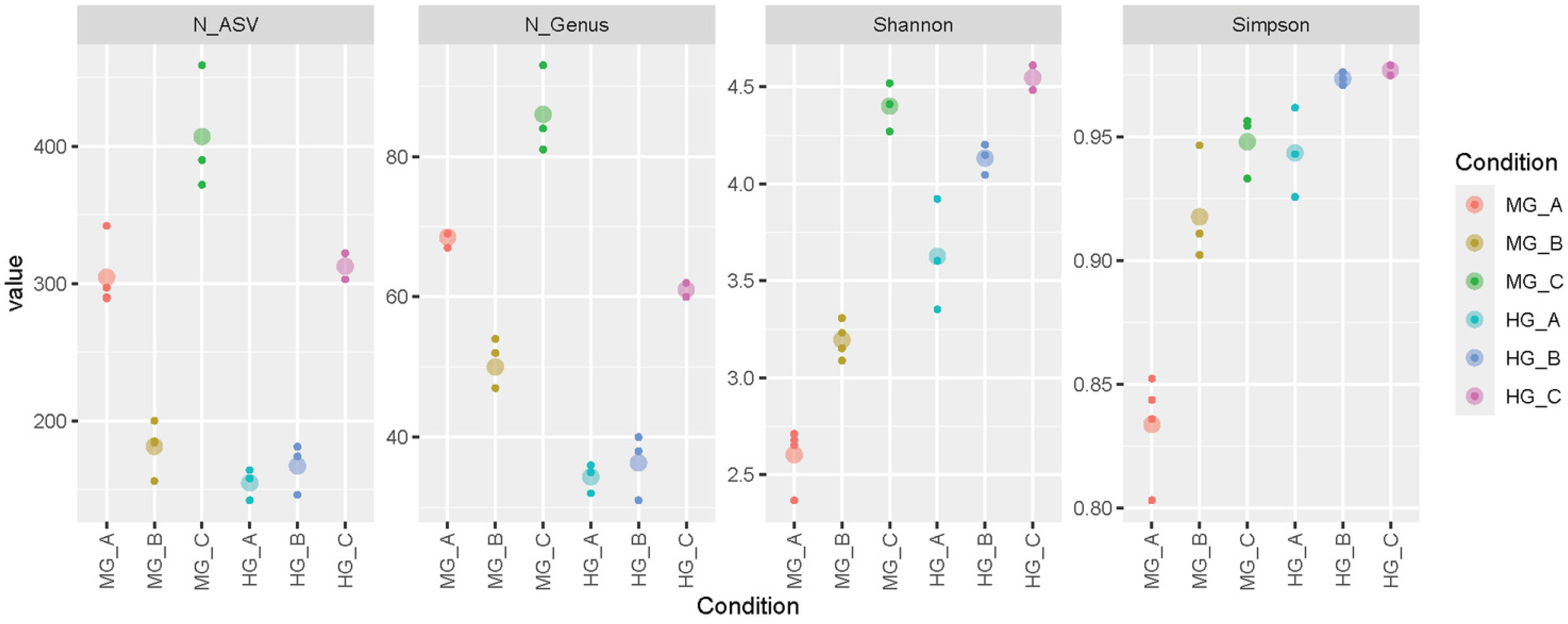
The number of ASVs, identified bacterial genera, and alpha diversity indices (i.e., Shannon and Simpson index values) per condition.

**Figure S2.**
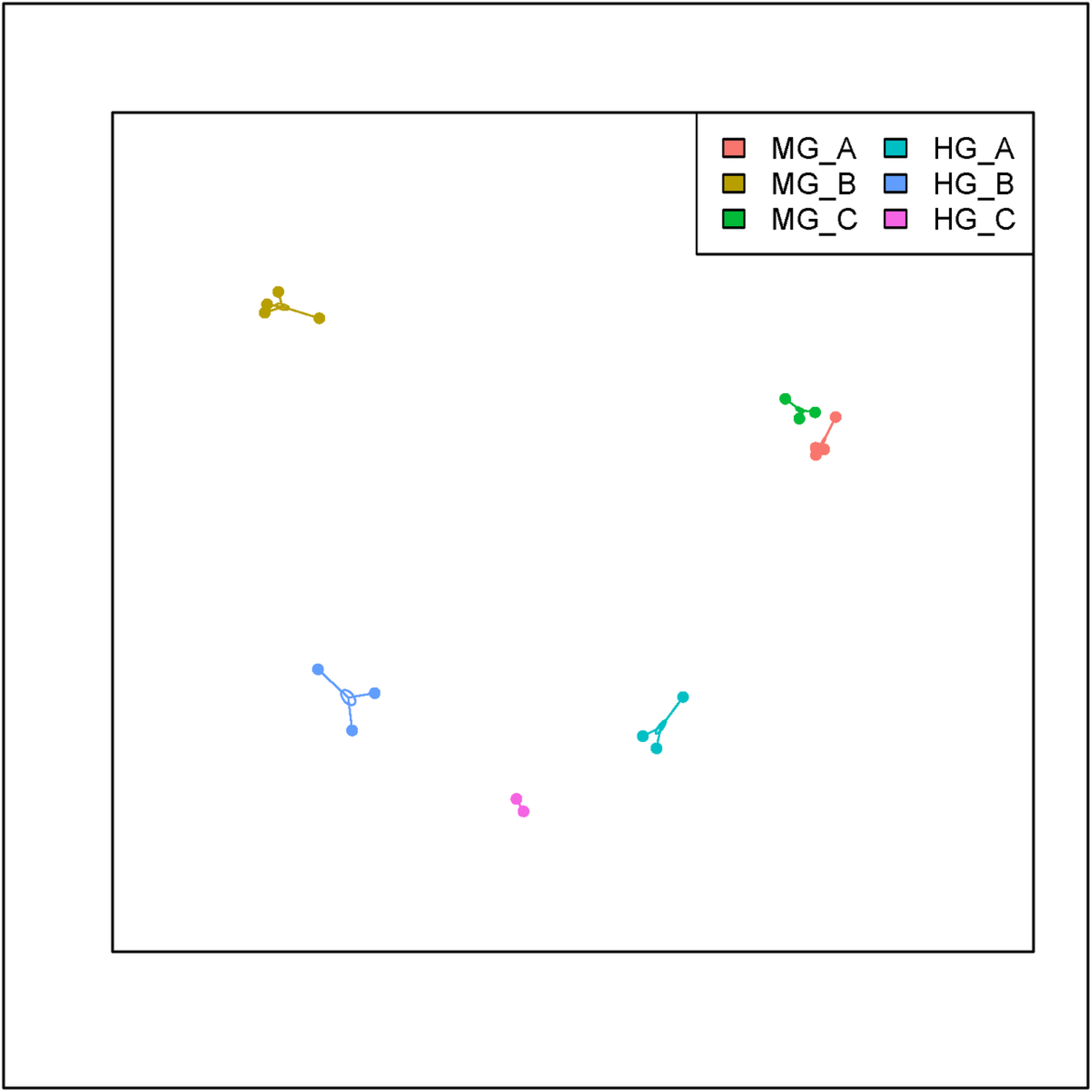
Beta-diversity analyses, as revealed by generalized principal coordinate analysis (PCoA), highlight the influence of diet and the gut compartment at the levels of ASV.

**Figure S3.**
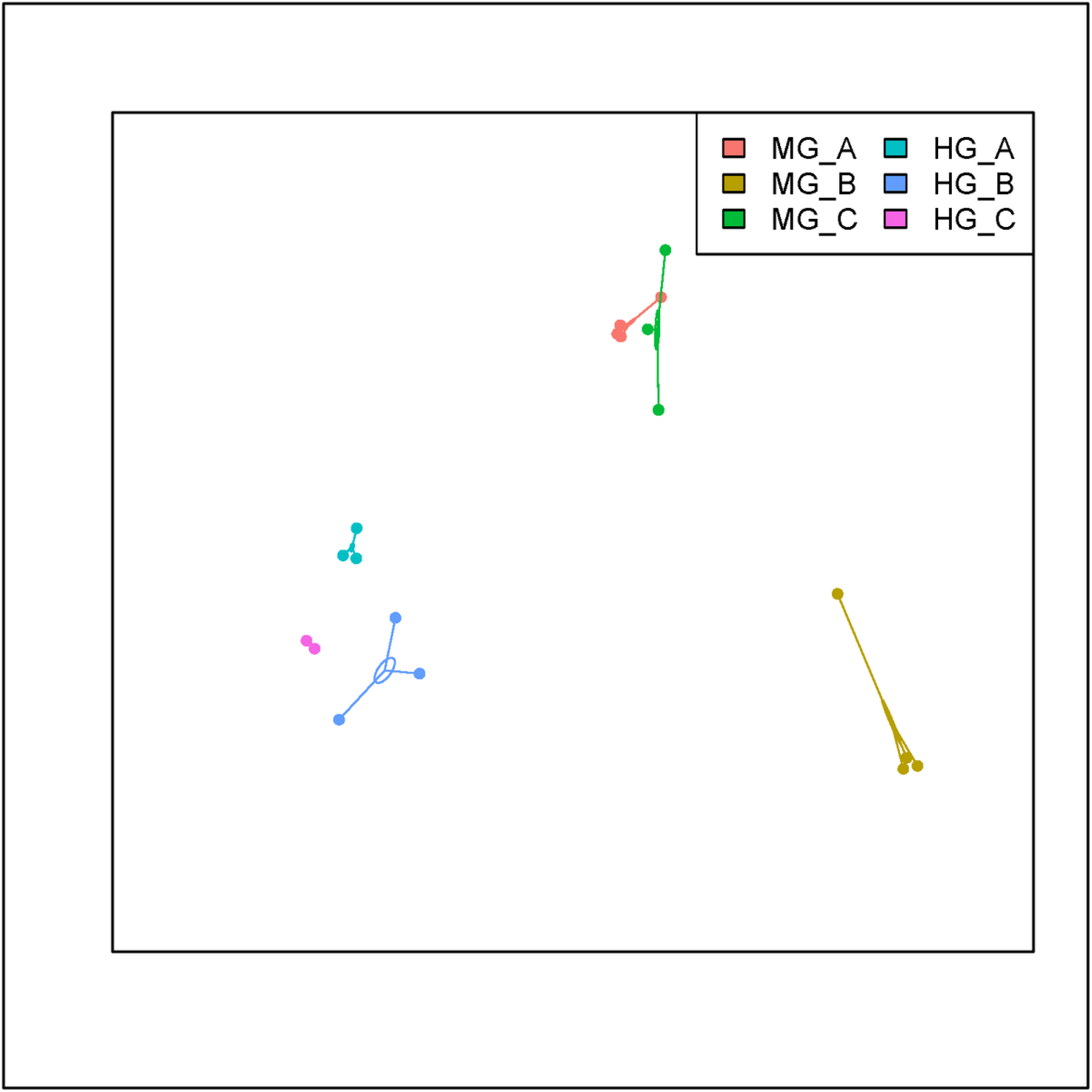
Beta-diversity analyses, as revealed by generalized principal coordinate analysis (PCoA), highlight the influence of diet and the gut compartment at the levels of phyla.

**Figure S4.**
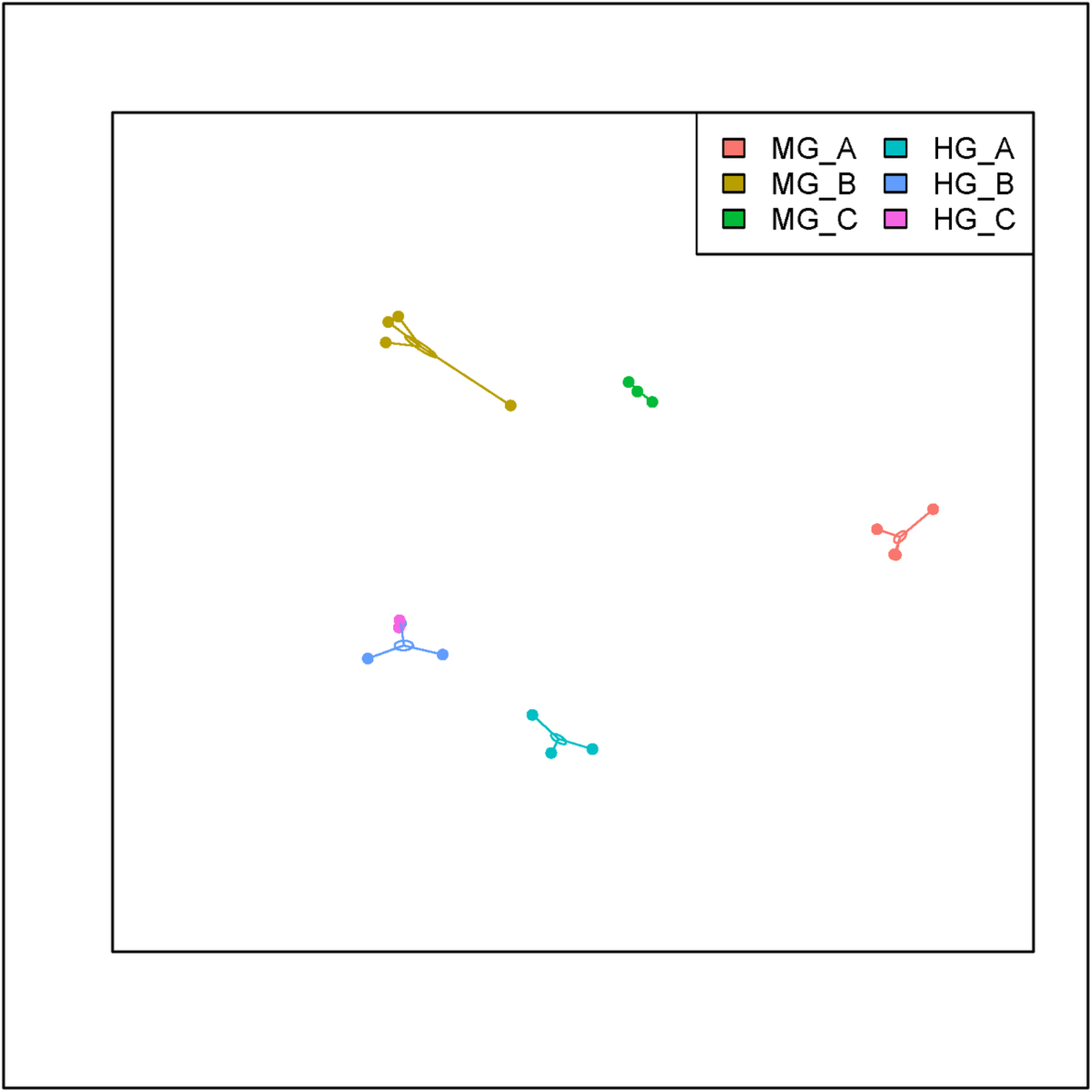
Beta-diversity analyses, as revealed by generalized principal coordinate analysis (PCoA), highlight the influence of diet and the gut compartment at the levels of families.

**Figure S5.**
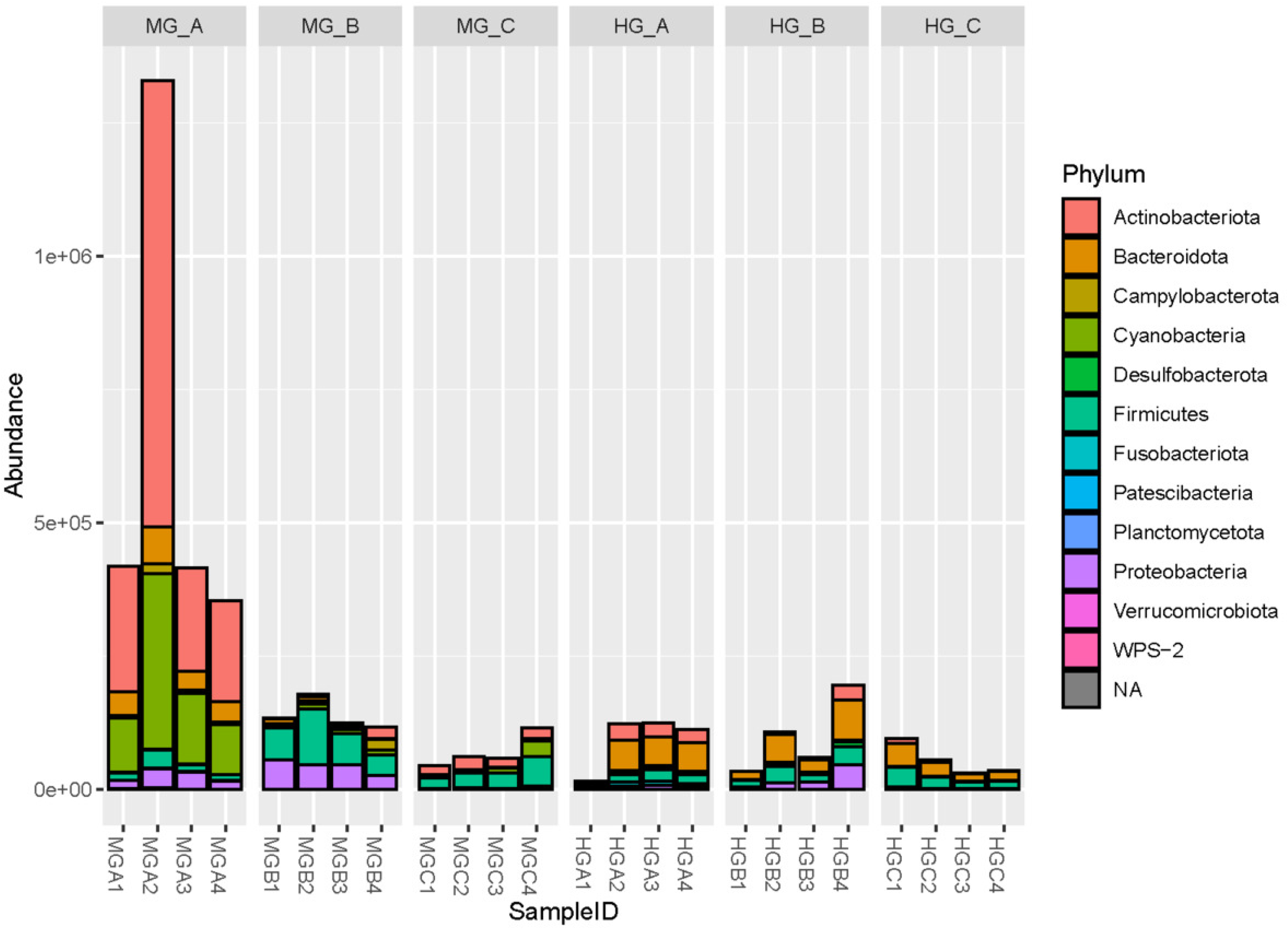
Absolute abundance (read counts) of bacterial phyla detected in the midgut (MG) and hindgut (HG) of BSFL fed on three distinct carbohydrate-rich diets: 100% fruit/vegetable waste (A), 100% potato scraps (B), and 100% forage (C). Metabarcoding analyses were performed on four biological replicates (1-4).

**Figure S6.**
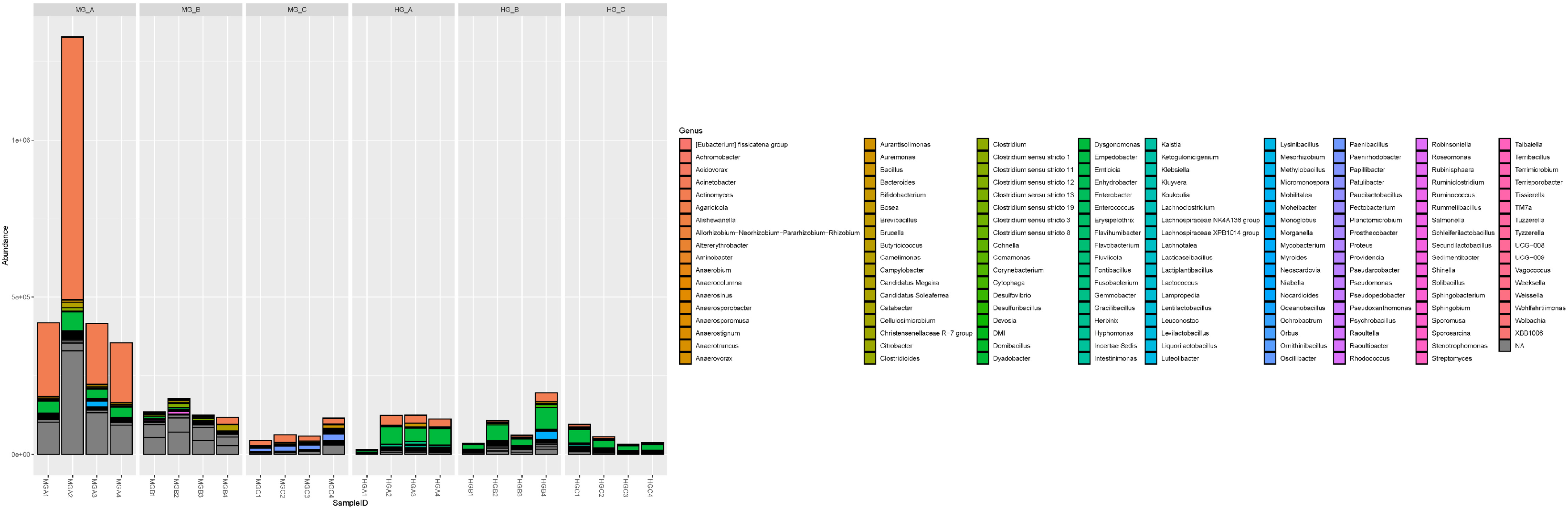
Absolute abundance (read counts) of bacterial genera (top 10) detected in the midgut (MG) and hindgut (HG) of BSFL fed on three distinct carbohydrate-rich diets: 100% fruit/vegetable waste (A), 100% potato scraps (B), and 100% forage (C). Metabarcoding analyses were performed on four biological replicates (1-4).

**Figure S7.**
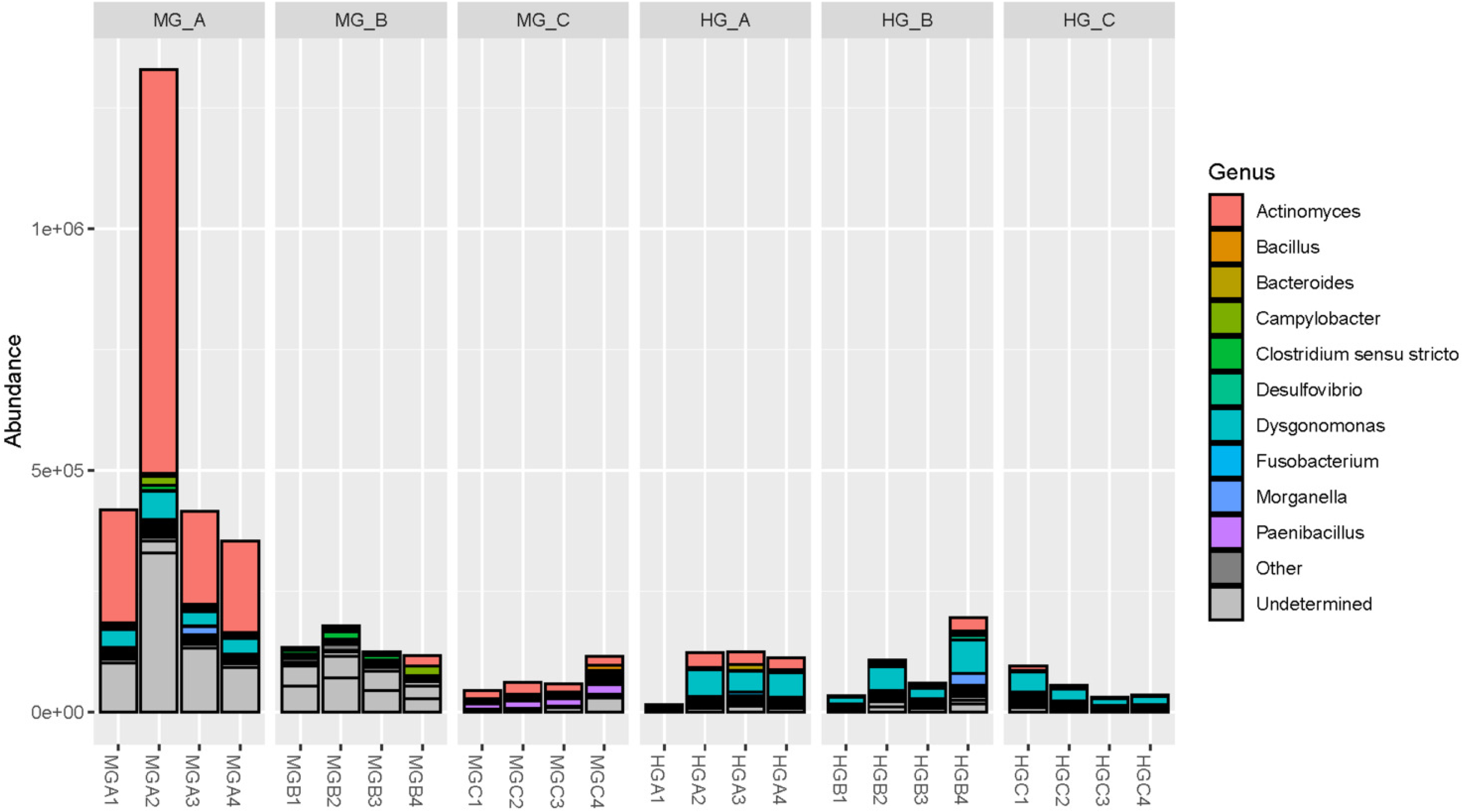
Absolute abundance (read counts) of the 10 most abundant bacterial genera detected in the midgut (MG) and hindgut (HG) of BSFL fed on three distinct carbohydrate-rich diets: 100% fruit/vegetable waste (A), 100% potato scraps (B), and 100% forage (C). Metabarcoding analyses were performed on four biological replicates (1-4).

**Figure S8.**
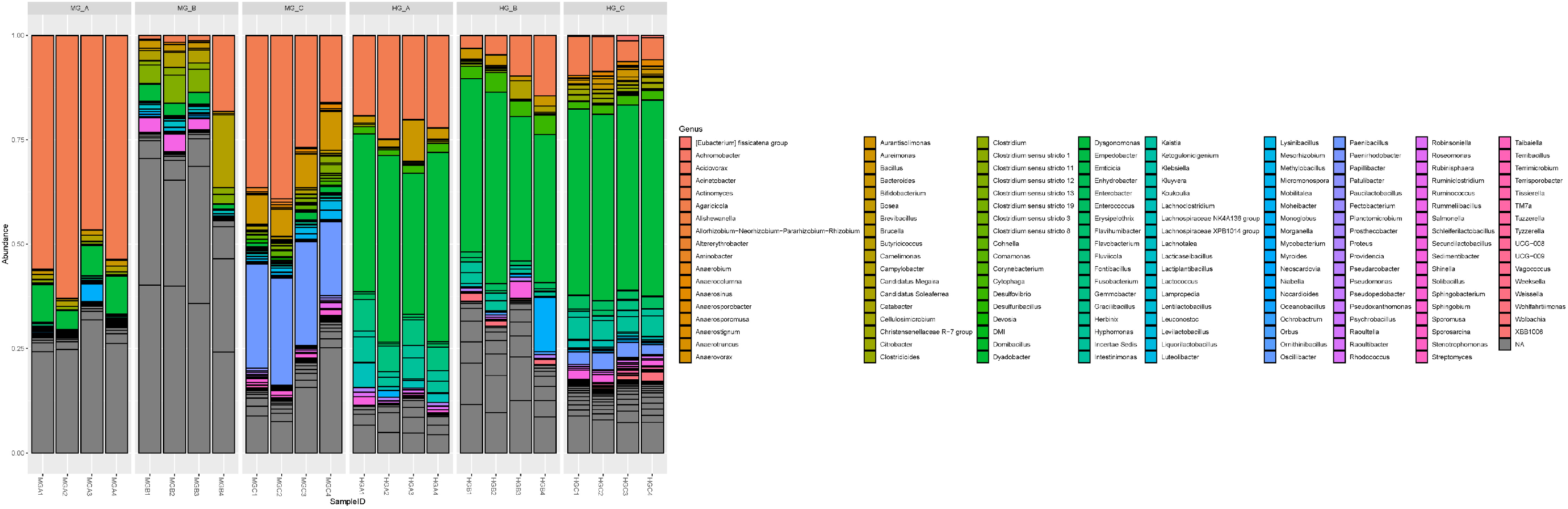
Relative abundance of the 10 most abundant bacterial genera in the midgut (MG) and hindgut (HG) of BSFL fed on three distinct carbohydrate-rich diets: 100% fruit/vegetable waste (A), 100% potato scraps (B), and 100% forage (C). Metabarcoding analyses were performed on four biological replicates (1-4).

**Figure S9.**
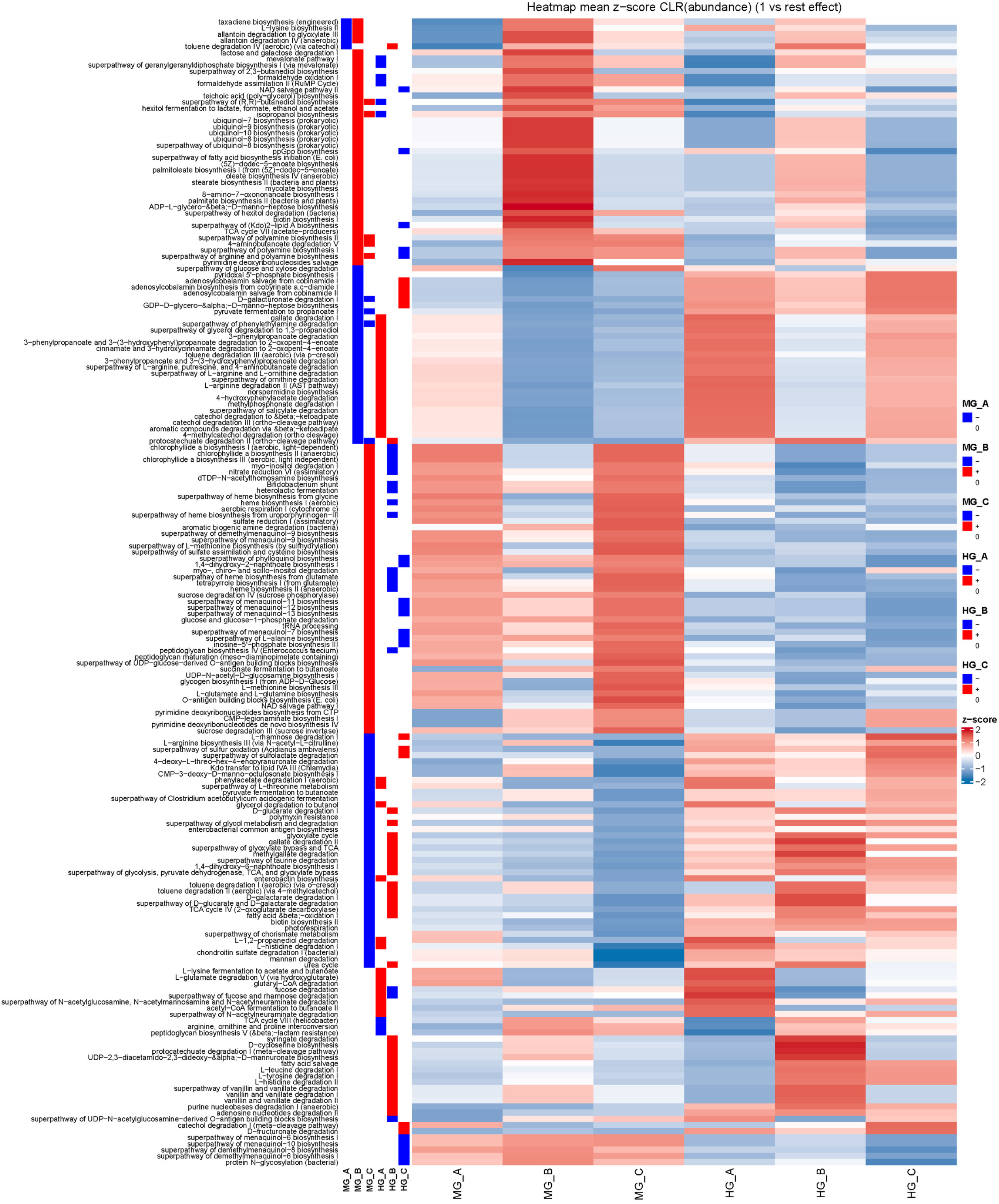
Predicted metabolic functions (PICRUSt2) differentially enriched across gut compartments and diets in *H. illucens*. Category abundances were centered log-ratio (CLR) transformed and standardized within each category as z-scores across sample groups. Red indicates a higher-than-average predicted relative abundance and blue a lower-than-average predicted relative abundance for a given functional category, whereas white indicates values close to the category mean. HG = Hind gut, MG = Mid gut, A corresponds to the fruit/vegetable waste diet, B to the potato scraps diet, and C to the forage diet.

